# The NLR helper protein NRC3 but not NRC1 is required for Pto-mediated cell death in *Nicotiana benthamiana*

**DOI:** 10.1101/015479

**Authors:** Chih-Hang Wu, Khaoula Belhaj, Tolga O. Bozkurt, Sophien Kamoun

**Affiliations:** The Sainsbury Laboratory, Norwich Research Park, Norwich, NR4 7UH, United Kingdom

## Abstract

Intracellular immune receptors of the nucleotide-binding leucine-rich repeat (NB-LRR or NLR) proteins often function in pairs, with “helper” proteins required for the activity of “sensors” that mediate pathogen recognition. The NLR helper NRC1 (<u>N</u>B-LRR protein <u>r</u>equired for HR-associated <u>c</u>ell death <u>1</u>) has been described as a signalling hub required for the cell death mediated by both cell surface and intracellular immune receptors in the model plant *Nicotiana benthamiana*. However, this work predates the availability of the *N. benthamiana* genome and whether NRC1 is indeed required for the reported phenotypes has not been confirmed. Here, we investigated the NRC family of solanaceous plants using a combination of genome annotation, phylogenetics, gene silencing and genetic complementation experiments. We discovered that a paralog of *NRC1*, we termed *NRC3*, is required for the hypersensitive cell death triggered by the disease resistance protein Pto but not Rx and Mi-1.2. NRC3 may also contribute to the hypersensitive cell death triggered by the receptor-like protein Cf-4. Our results highlight the importance of applying genetic complementation to validate gene function in RNA silencing experiments.

## INTRODUCTION

Plants defend against pathogens using both cell surface and intracellular immune receptors (Dodds & Rathjen, 2010; Win *et al*., 2012). Plant cell surface receptors include receptor-like kinases (RLKs) and receptor-like proteins (RLPs), which respond to pathogen-derived apoplastic molecules (Boller & Felix, 2009; Thomma *et al*., 2011). In contrast, plant intracellular immune receptors are typically nucleotide-binding leucine-rich repeat (NB-LRR or NLR) proteins, which respond to translocated effectors from a diversity of pathogens (Eitas & Dangl, 2010; Bonardi *et al*., 2012). These receptors engage in microbial perception either by direct binding pathogen molecules or indirectly by sensing pathogen-induced perturbations (Win *et al*., 2012). However, signaling events downstream of pathogen recognition remain poorly understood.

In addition to their role in microbial recognition, some NLR proteins contribute to signal transduction and/or amplification (Gabriels *et al*., 2007; Bonardi *et al*., 2011; Cesari *et al*., 2014). An emerging model is that NLR proteins often function in pairs, with “helper” proteins required for the activity of “sensors” that mediate pathogen recognition (Bonardi *et al*., 2011; Bonardi *et al*., 2012). Among previously reported NLR helpers, NRC1 (<u>N</u>B-LRR protein <u>r</u>equired for HR-associated <u>c</u>ell death <u>1</u>) stands out for having been reported as a signalling hub required for the cell death mediated by both cell surface immune receptors such as Cf-4, Cf-9, Ve1, and LeEix2, as well as intracellular immune receptors, namely Pto, Rx, and Mi-1.2 (Gabriels *et al*., 2006; Gabriels *et al*., 2007; Sueldo, 2014). However, most of these studies predate the availability of the *N. benthamiana* genome and it remains questionable whether NRC1 is indeed required for the reported phenotypes.

Functional analyses of NRC1 were performed using virus-induced gene silencing (VIGS) (Gabriels *et al*., 2007), a method that is popular for genetic analysis in several plant systems, particularly the model solanaceous plant *Nicotiana benthamiana* (Burch-Smith *et al*., 2004). However, interpretation of VIGS can be problematic as the experiment can result in off-targets silencing (Senthil-Kumar & Mysore, 2011). In addition, heterologous gene fragments from other species (e.g. tomato) have been frequently used to silence homologs in *N. benthamiana*, particularly in studies that predate the sequencing of the *N. benthamiana* genome (Burton *et al*., 2000; Liu *et al*., 2002b; Lee *et al*., 2003; Gabriels *et al*., 2006; Gabriels *et al*., 2007; Senthil-Kumar *et al*., 2007; Oh *et al*., 2010). In the NRC1 study, a fragment of a tomato gene corresponding to the LRR domain was used for silencing in *N. benthamiana* (Gabriels *et al*., 2007). Given that a draft genome sequence of *N. benthamiana* has been generated (Bombarely *et al*., 2012) and silencing prediction tools have become available (Fernandez-Pozo *et al*., 2014), we can now design better VIGS experiments and revisit previously published studies.

Two questions arise about the NRC1 study. First, is there a *NRC1* ortholog in *N. benthamiana*? Second, has the gene silencing with the tomato *NRC1* fragment affected other *N. benthamiana* genes with similarity to *NRC1*? In this study, we investigated the NRC family of solanaceous plants using a combination of genome annotation, phylogenetics, gene silencing and genetic complementation experiments. We discovered that a paralog of *NRC1*, we termed *NRC3*, is required for the hypersensitive cell death triggered by Pto but not Rx and Mi-1.2. NRC3 may also contribute to the hypersensitive cell death triggered by Cf-4. Our results highlight the importance of applying genetic complementation to validate gene function in RNA silencing experiments.

## RESULTS AND DISCUSSION

### NRC1 and related NLR proteins form a complex family in solanaceous plants

To identify putative homologs of NRC1 in *N. benthamiana*, potato, and tomato genomes, we performed a BLASTP (Altschul *et al*., 1990) search against the predicted protein databases in Sol Genomics Network (SGN) using the sequence of tomato NRC1 (Solyc01g090430) as a query. Protein hits with more than 50% identity to tomato NRC1 were selected for further analysis. Phylogenetic analysis and sequence comparisons indicated that the NRC family is composed of three subclades (NRC1-3). Surprisingly, a *N. benthamiana* ortholog was missing in the NRC1 subclade and a tomato ortholog was also missing in the NRC3 subclade.

To determine whether the missing sequences are due to misannotations in the tomato and *N. benthamina* genomes, we searched all the available nucleotide and protein databases of *N. benthamiana* and tomato in SGN with representative NRC sequences. We failed to identify sequences that show high similarity to NRC1 in *N. benthamiana*, even after blast searches against scaffolds and contigs sequences in both SGN and the *Nicotiana benthamiana* genome database (www.benthgenome.com) (Naim *et al*., 2012; Nakasugi *et al*., 2014). We, therefore, concluded that NRC1 is probably missing in the *N. benthamiana* genome although it may have been somehow omitted from the assembly. In contrast, using TBLASTN searches, we detected a misannotated tomato gene in contig SL2.40ct02653 with high similarity to potato NRC3. Based on sequence comparisons, this gene has three exons and two introns; the first two exons were annotated as Solyc05g009630 whereas the third exon was missing in the annotation (Fig. S1a). To validate the sequence and expression of tomato *NRC3*, we designed primers based on our predicted full-length sequence and performed PCR using tomato cDNA and genomic DNA as template. We successfully amplified a fragment from genomic DNA and cDNA (Fig. S1b). The amplified cDNA fragment was cloned and sequenced. The identity between tomato NRC3 and potato NRC3 is 95%, consistent with our interpretation that the encoding gene is the *NRC3* ortholog in tomato (Fig. S1c, Fig. S2).

Phylogenetic analyses that include the newly identified tomato NRC3 revealed that the sequences in the NRC family fall into three subclades that are supported by robust bootstrap values (Fig. 1). According to the genome information of potato and tomato, sequences in these three clades are located on three different chromosomes (Fig. 1), consistent with the view that genes in the same NRC subclade are orthologous.

**Figure 1.**
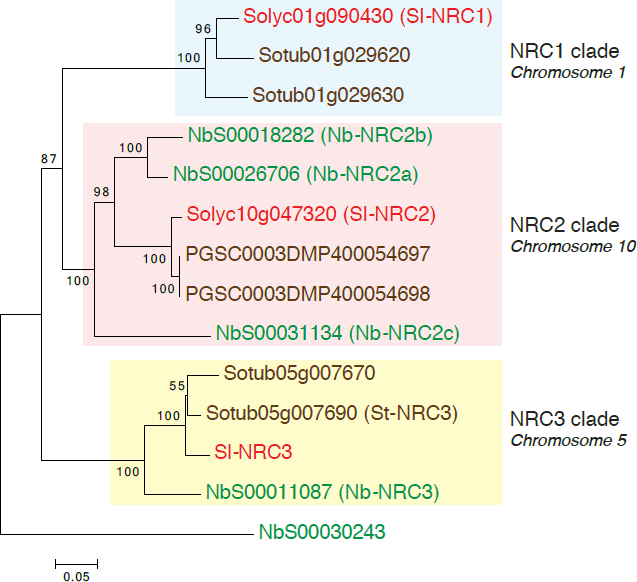
Phylogenetic tree of NRC homologs in solanaceous plants. Sequences with more then 50% identity to tomato NRC1 (Sl-NRC1) were analysed in MEGA6 to generate Neighbor-joining tree. Chromosome assignments are based on potato and tomato genome. Numbers at branches indicate bootstrap support (1000 replicates), and scale indicates the evolutionary distance in substitution per nucleotide. Sequences from tomato, potato and *N. benthamiana* are marked in red, brown and green, respectively.

### Silencing of NRC family members suppresses cell death mediated by Pto

We exploited the *N. benthamiana* genome sequence and associated gene silencing target prediction tool (SGN VIGS tool; http:// http://vigs.solgenomics.net) to analyse the specificity of the *NRC1* VIGS fragment that was used in the *NRC1* VIGS experiments (Gabriels *et al*., 2007). We found that this Sl*NRC1* fragment, which matches the LRR domain, would most probably target the *N. benthamiana* genes *NbNRC2a/b* and *NbNRC2c*, and possibly *NbNRC3*. This prompted us to test the degree to which silencing of the individual *NRC2a/b*, *NRC2c* or *NRC3* genes could suppress the cell death mediated by different immune receptors. To design more specific constructs for silencing individual *NbNRC* paralogs, we analysed the *NbNRC* sequences with the VIGS tool. The 5’ coding regions of each gene provided the highest specificity and were selected to design new gene silencing constructs. *N. benthamiana* plants were subjected to VIGS and challenged with the immune receptors Pto, Cf-4, Rx and Mi-1.2. Silencing of *NRC2a/b* or *NRC3* moderately but significantly reduced the cell death mediated by Pto and Cf-4 but not Rx and Mi-1.2 (Fig. 2a). In contrast, silencing of *NRC2c* resulted in a minor reduction in Cf-4 cell death but had no significant effect on Pto. Semi-quantitative RT-PCR indicated that the VIGS constructs reduced the expression of the targeted gene with no detectable effects on the other paralogs (Fig. 2b).

**Figure 2.**
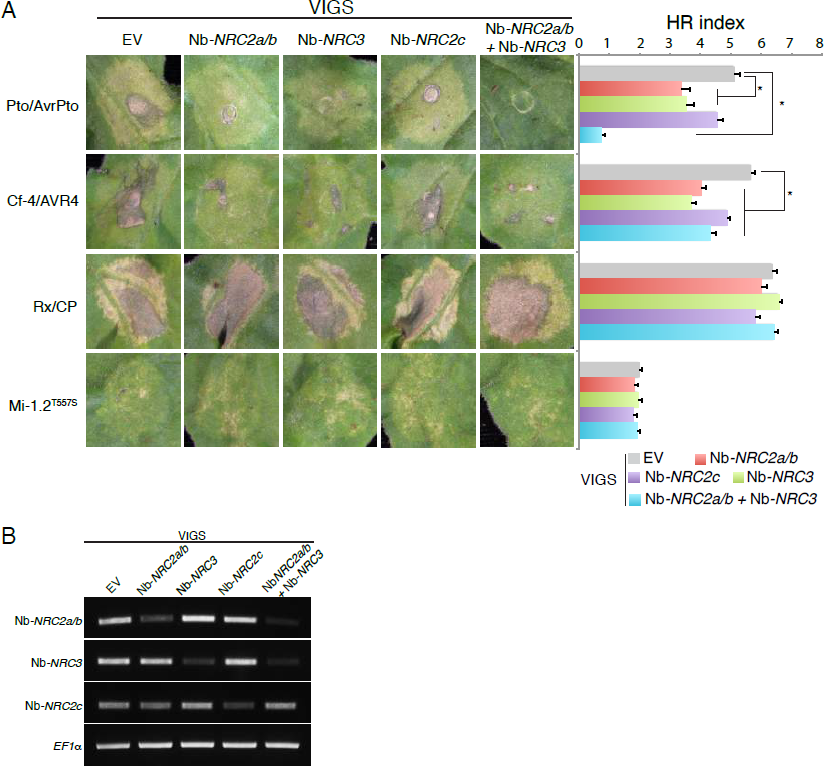
Silencing of *N. benthamiana NRC* homologs suppress cell death mediated by Pto/AvrPto. **(A)** Silencing (VIGS) of Nb-*NRC* homologs suppress Pto-medaited cell death. Immune receptors and corresponding AVR proteins, or autoactive immune receptor (Mi-1.2^T557S^), were transiently expressed in *N. benthamiana* leaves with different silencing constructs of *NRC* homologs. The HR results are presented with representative images. HR index was established at 7 days post infiltration (dpi). Bars represent mean ± SE of 24 infiltrations from one biological repeat. Statistical differences among the samples were analysed by Student’s *t*-test (\**p*-value < 0.01). Experiments were performed at least three times with similar results. **(B)** Semi-quantitative RT-PCR of *NRC*-silenced *N. benthamiana* leaves. Leaves were collected three weeks after virus inoculation. Elongation factor-1α (*EF1α*) was used as an internal control.

Next, we combined the two *NRC2a/b* and *NRC3* fragments in one construct with the aim of obtaining more robust phenotypes. Interestingly, the double-silencing construct that targets both *NRC2a/b* and *NRC3* dramatically suppressed Pto-mediated cell death close to background levels (Fig. 2a,b), suggesting that *NRC2a/b* and *NRC3* may be functional redundant in mediating Pto-mediated cell death. In contrast, the intensity of Cf-4-mediated cell death remained unchanged relative to the individual silencing of *NRC2a/b* and *NRC3*. As with the individual silencing constructs, Rx and Mi-1.2-mediated cell death were unaffected. These results clearly show that silencing of *NRC2a/b* and *NRC3* affects the cell death induced by Pto and probably Cf-4, but does not alter Rx- and Mi-mediated cell death.

### NRC3 mediates Pto-induced cell death

To clarify which of NRC2a/b and NRC3 are implicated in Pto-mediated cell death, we performed complementation experiments to determine which of the tomato *NRC* genes can rescue Pto-mediated cell death when the *N. benthamiana* endogenous *NRC* genes are silenced (Fig. 1 and Fig. 2). Our experiment was motivated by the observation that the tomato NRC sequences are probably divergent enough from the *N. benthamiana* ones to be resilient to silencing. These experiments revealed that *SlNRC3* partially rescued Pto-elicited cell death, *SlNRC2* showed weak complementation activity but *SlNRC1* did not (Supplemental Fig. 3a). Based on these results, we conclude that, unlike *Sl*NRC3, *Sl*NRC1 and *Sl*NRC2 are unlikely to be fully required for Pto-elicited hypersensitive response.

**Figure 3.**
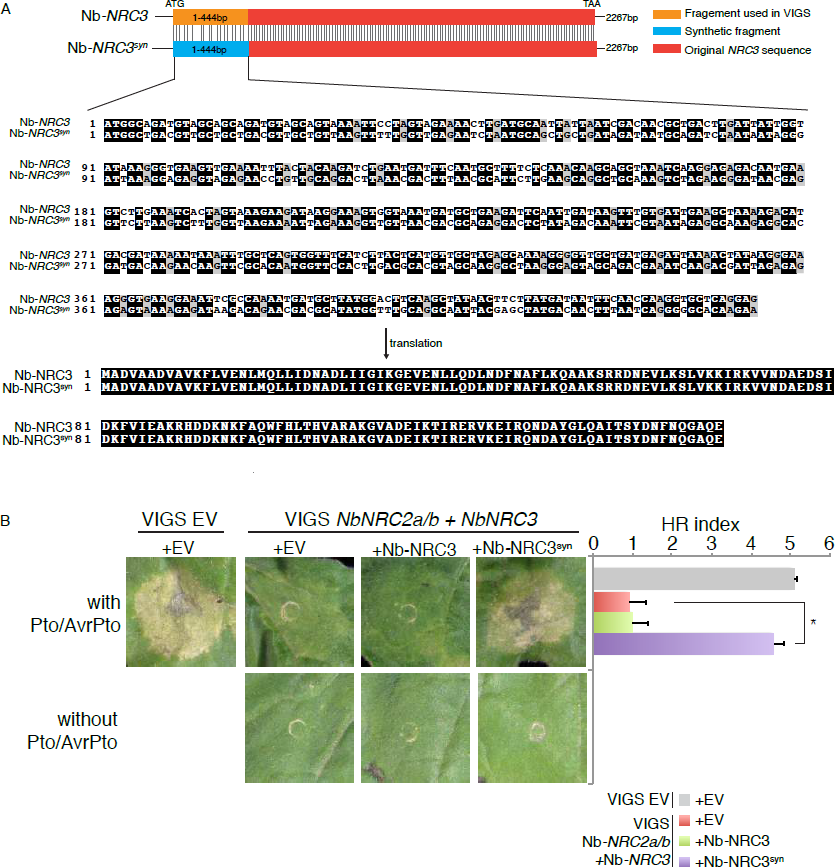
Synthetic Nb-NRC3 rescues Pto-mediated cell death in *NRC*-silenced *N. benthamiana*. **(A)** Schematic representation of experimental design, and DNA and protein sequences of the synthetic region. Shuffled synonymous codons were introduced in the synthetic sequence (*Nb-NRC3^syn^*) without changing the identity in protein sequence. **(B)** Nb-NRC3^syn^ rescues the cell death of Pto/AvrPto. Nb-NRC3 and Nb-NRC3^syn^ were co-expressed with Pto/AvrPto in *NRC*-silenced *N. benthamiana* leaves. Expression of Pto/AvrPto in VIGS empty vector (EV) and expression of Nb-NRC3 and Nb-NRC3^syn^ without Pto/AvrPto were used as control. HR index was established at 7 days post infiltration (dpi). Bars represent mean ± SE of 14 infiltrations from one biological repeat. Statistical differences among the samples were analysed by Student’s *t*-test (\**p*-value < 0.01). Experiments were performed at least three times with similar results.

To challenge this finding, we tested the extent to which overexpression of the tomato *NRC* genes enhances the hypersensitive cell death mediated by different immune receptors. Of the three genes we tested, *SlNRC3* was the only one to enhance Pto-mediated cell death (Fig. S3b). Overexpression of *SlNRC3* also enhanced Cf-4-mediated cell death, consistent with our previous conclusion that NRC3 may contribute to Cf-4 signalling. Taken together, the results of the complementation and overexpression experiments indicate that NRC3, but not NRC1 nor NRC2, mediates signalling in *N. benthamiana* following activation of the Pto/Prf intracellular receptors and Cf-4 cell surface receptor-like protein. However, it remains possible that the failure of NRC1 and NRC2 to rescue and enhance cell death is due to problems related to heterologous expression.

### Genetic complementation with a synthetic *NbNRC3* gene

We aimed to confirm that the *N. benthamiana NRC3* gene is the causal gene implicated in the cell death elicited by Pto given that the complementation assays were performed with tomato *NRC* genes. To achieve this, we generated a synthetic version of *NbNRC3*, termed *NbNRC3^syn^*, with shuffled synonymous codon sequences and that should be divergent enough to evade VIGS (Fig. 3a). Expression of *NbNRC3^syn^* in *NRC2*- and *NRC3*-silenced *N. benthamiana* leaves rescued the cell death mediated by Pto, whereas the endogenous *NbNRC3* gene failed to complement (Fig. 3b). These experiments clearly demonstrate that *Nb*NRC3 is the causal *N. benthamiana* protein that contributes to hypersensitive death following Pto perception of AvrPto.

## CONCLUSIONS

In summary, we revisited the role of NRC1 as a helper NLR protein and discovered that NRC3 rather than NRC1 is the causal protein required for Pto-mediated cell death in *N. benthamiana*. Therefore, the previously proposed model of NRC1 as a signaling hub for multiple immune receptors postulated by Gabriels *et al*. (2007) needs to be revised. In fact, the *N. benthamiana* genome appears to lack an ortholog of tomato *NRC1* (Fig. 1). Furthermore, although NRC3 is required for the hypersensitive cell death induced by Pto, it doesn’t seem to be required for the response elicited by Rx and Mi-1.2. The previous finding of Gabriels *et al*. (2007) that silencing of *NRC1* suppresses Rx and Mi-1.2 –mediated cell death may be due to an effect on other NRC1-like sequences in *N. benthamiana*. We did, however, observe that *NRC3* silencing partially reduced the cell death induced by Cf-4 as reported earlier (Gabriels *et al*., 2007). However, the cell death elicited by Cf-4 is relatively weak and the effect of NRC3 silencing is not as dramatic as in the case of Pto-mediated cell death (Fig. 2).

Our findings emphasize the importance of following RNA silencing experiments with genetic complementation assays to minimize the risk of misinterpretation due to off-target effects (Kumar *et al*., 2006; Jonchere & Bennett, 2013; Pliego *et al*., 2013). Genetic complementation can be performed using genes from a different species or using a silencing-resilient synthetic version of the gene with shuffled codon sequences. We recommend that genetic complementation be applied to RNA silencing experiment whenever possible to avoid gene misidentification.

## MATERIALS AND METHODS

### Identification of NRC homologs and phylogenetic analyses

Tomato NRC1 (Solyc01g090430) was used for searching homologs in the predicted protein databases (*N. benthamiana* Genome v0.4.4 predicted protein, Tomato proteins ITAG release 2.30, and Potato ITAG release 1 predicted proteins) on Solanaceae Genomics Network (SGN). Sequences with more than 50% identity were collected for further analysis. NRC2 homologs in potato were missing in Potato ITAG release 1 predicted proteins database. Therefore, two NRC2 sequences of potato identified in Potato PGSC DM v3.4 protein sequences were included in the analysis. The phylogenetic tree of NRC homologs was built using MEGA6-Beta2 (Tamura *et al*., 2013) with Neighbor-joining method with bootstrap values based on 1000 iterations. Chromosome assignments of NRC homologs were based on tomato and potato genomes.

### Cloning of solanaceous NRC homologs

Cloning of tomato *NRC* homologs was performed with the Gateway cloning kit following the manufacturer’s instruction (Invitrogen). Primer pairs used in the cloning are listed as follows: *S*lNRC1-F/R (5’-CACCATGGTTGATGTAGGGG-TTGAATTTC-3’ and 5’-CTAAGAAGCTGTCTGTACATCAGAATC-3’), *Sl*NRC2-R/F (5’-CACCATGGCGAACGTAGCAGTGGAATTTC-3’ and 5’-TCAGAGATCAGGAGGGAATATGGAAAG-3’), and *Sl*NRC3-F/R (5’-CACCATGGCGGATGTAGCAGTAAAGTTCTTA-3’ and 5’-TTACAATCCAAG-ATCATGAGGGAAT-3’). The amplified fragments from tomato cDNA were cloned into pENTR/D-TOPO (Invitrogen) and then introduced into the pK7WG2 destination vector (Karimi *et al*., 2002) by Gateway LR recombination enzyme (Invitrogen). *N. benthamiana NRC3* was amplified with the corresponding primer pairs (5’- AATTGGTCTCTAATGGCAGATGCAGTA-GTGAATTTTCTGGTG-3’ and 5’- ATTGGTCTCGAAGCTTACTGTGTGGCC-TTGGATCCAGCTTC-3’) from cDNA and cloned into pCR8/GW/TOPO (Invitrogen) by TA cloning. The fragment was then subcloned into pICH86988 with Golden Gate cloning (Weber *et al*., 2011). The synthetic fragment of *NbNRC3* was designed manually to introduce synonymous substitution in almost every codon, which reduces the nucleotide sequence identity to 64% and no identical fragment longer than 5bp. Gene synthesis was performed with GENEWIZ. Inc. The synthetic fragment was then subcloned into pICH86988 together with the rest part of *NbNRC3* to generate full length *NbNRC3^syn^*.

### Virus-induced gene silencing (VIGS)

VIGS was performed in *N. benthamiana* as described by Liu et al (2002a). Suspensions of *Agrobacterium tumefaciens* strain GV3101 harbouring TRV RNA1 (pYL155) and TRV RNA2 (pYL279) (Liu *et al*., 2002a) with corresponding fragments from indicated genes were mixed in a 2:1 ratio in infiltration buffer (10 mM MES, 10mM MgCl_2_, and 150 mM acetosyringone, pH5.6) to a final OD_600_ of 0.3. Two-week-old *N. benthamiana* plants were infiltrated with the agrobacteria for VIGS assays, and systemic leaves were used three weeks later for further agroinfiltrations. For silencing of *NRC* homologs in *N. benthamiana*, 5’ coding region of each gene (*NbNRC2a/b*, 1-429b; *NbNRC2c*, 1-426bp; *NbNRC3*, 1-444bp) were cloned into TRV RNA2 vector. The fragments of *NbNRC2a/b* and *NbNRC3* were then fused by overlay PCR and cloned into TRV RNA2 vector to generate a construct which silence both genes simultaneously.

### Transient gene expression and complementation assays

*In planta* transient agroinfiltration assays were performed according to methods described previously (Bos *et al*., 2006). *A. tumefaciens* strains carrying the expression constructs were adjusted to the indicated concentration in the infiltration buffer (10 mM MES, 10 mM MgCl_2_, and 150 mM acetosyringone, pH5.6). Five-week-old *N. benthamiana* plants (i.e. three weeks after virus inoculation) were used for cell death tests in VIGSed background whereas four-week-old *N. benthamiana* plants were used for enhanced cell death assays. The final concentration of agrobacteria (OD_600_) in most cell death assays (Fig. 2, 3 and Fig. S3a) are indicated as follows: Pto, 0.6 (de Vries *et al*., 2006); AvrPto, 0.1 (de Vries *et al*., 2006); Cf-4, 0.5 (Liebrand *et al*., 2012), AVR4, 0.5 (Van der Hoorn *et al*., 2000); Rx, 0.2 (Tameling & Baulcombe, 2007); CP, 0.1 (Tameling & Baulcombe, 2007); Mi-1.2^T557S^, 0.8 (Lukasik-Shreepaathy *et al*., 2012). For enhanced cell death assays (Fig. S3b), lower concentrations of agrobacteria were used to reduce the cell death background. The concentration (OD_600_) is indicted as follow: Pto (0.4), AvrPto (0.03), Cf-4 (0.3), AVR4 (0.4), *Sl*NRC1 (0.2), *Sl*NRC2 (0.2), and *Sl*NRC3 (0.2). The hypersensitive response (HR) cell death phenotype was scored at 7 dpi, according to an arbitrary scale from 0 (no HR observed) to 7 (confluent necrosis) modified from Segretin et al. (Segretin *et al*., 2014).

### RNA extraction and semi-quantitative RT-PCR

Plant total RNA was extracted using RNAeasy Mini Kit (Qiagen). 2mg RNA of each sample was subject to first strand cDNA synthesis using Ominiscript RT Kit. Semi-quantitative RT-PCR was performed using DreamTaq (Thermo Scientific) with 25 to 30 amplification cycles followed by electrophoresis with 2% agarose gel stained with Ethidium bromide. Primer pairs used in the PCR reaction are listed as follows: *NbNRC2a/b-RT-F/R* (5’-AGTGGATGAGA-GTGTGGGTG-3’ and 5’-AAGCAGGGATCTCAAAGCCT-3’), *NbNRC2c-RT-R/F* (5’-TCAAAACATGCCGTGTTCAT-3’ and 5’-CCTGCGGGTTTTGTACT-GAT-3’) and *NbNRC3-RT-F/R* (5’- CCTCGAAAAGCTGAAGTTGG-3’ and 5’-TGTCCCCTAAACGCATTTTC-3’). Primers for internal control *NbEF1a* was described previously (Segonzac *et al*., 2011).

### Nomenclature

The nomenclatures of the genes mentioned in this article are based on the orthology to NRC1 (<u>N</u>B-LRR protein <u>r</u>equired for HR-associated <u>c</u>ell death <u>1</u>) and results of phylogenetic analysis as shown in Fig. 1. Species names are indicated as two-letter prefixes to the gene names; *Sl* is used for tomato (*Solanum lycopersicum*), *St* is used for potato (*Solanum tuberosum*) and *Nb* is used for *Nicotiana benthamiana*.

## ACKNOWLEDGEMENTS

We thank Florian Jupe for discussion about identification of NRC homologs in solanaceous plants. This project was funded by the Biotechnology and Biological Science Research Council (BBSRC), the European Research Council (ERC), and the Gatsby Charitable Foundation.

## Supplemental information

**Supplemental Figure 1.**
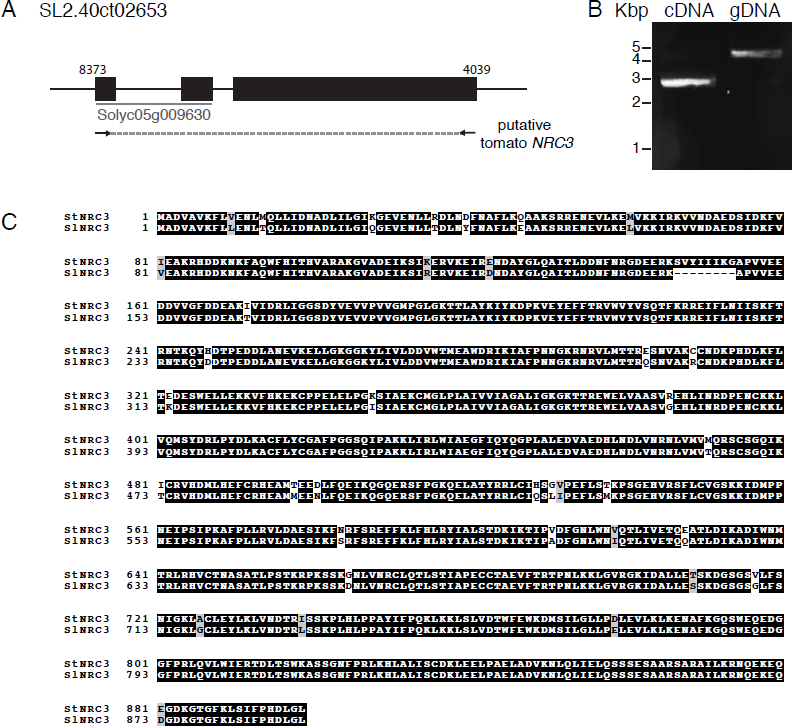
Cloning of tomato NRC3. **(A)** Schematic representation of predicted gene model of tomato *NRC3* (Sl-*NRC3*). Black boxes represent the three exons of Sl-*NRC3*. Numbers on the top indicate position of the start and stop codon in the contig. The first two exons were annotated as Solyco05g009630 in the SGN database. **(B)** PCR amplification of Sl-*NRC3* with tomato cDNA and genomic DNA. **(C)** Sequence alignment of tomato NRC3 with potato NRC3 (Sotub05g007690). Sequences were aligned with ClustalW2 and analysed by BoxShade. Identical amino acids are highlighted in black and conserved amino acids in grey

**Supplemental Figure 2.**
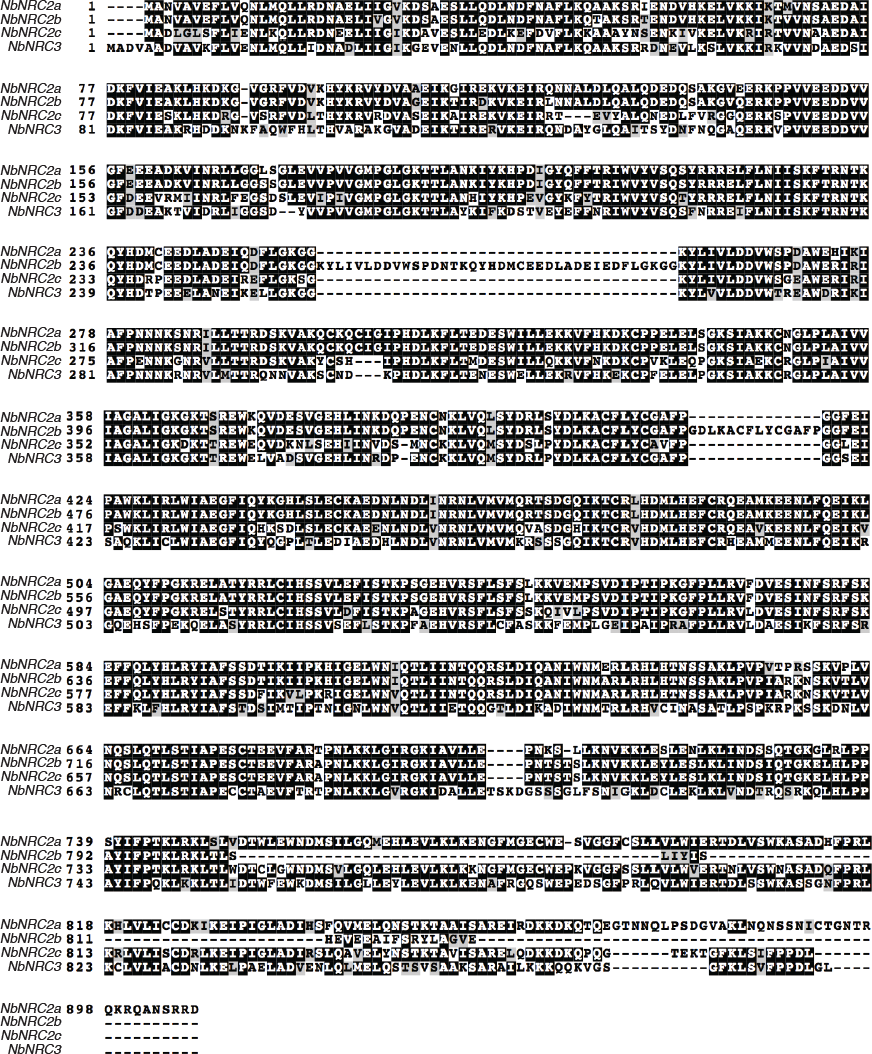
Protein sequence alignment of NRC homologs in *N. benthamiana*. Predicted sequences of NRC homologs in *N. benthamiana* were aligned with ClustalW2 and analysed by BoxShade. Identical amino acids are highlighted in black and conserved amino acids in grey

**Supplemental Figure 3.**
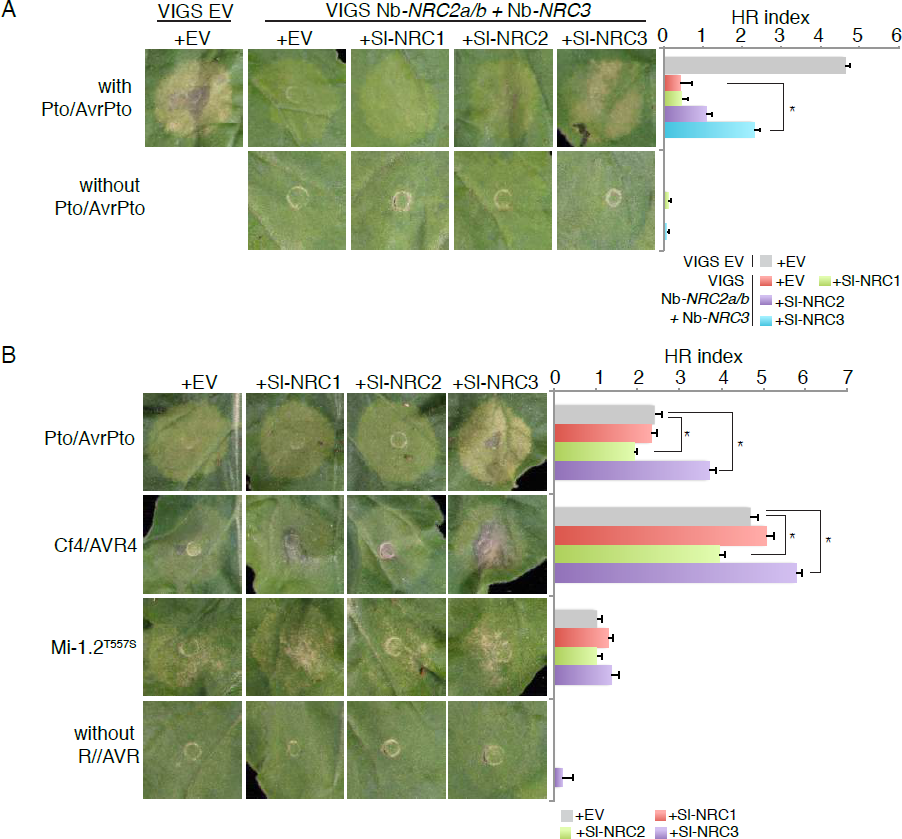
Tomato NRC3 but not NRC1 or NRC2 is required for Pto-mediated cell death in *N. benthamiana*. **(A)** Complementation assay with tomato NRC homologs. Tomato NRC homologs (Sl-NRC1, Sl-NRC2 and Sl-NRC3) were co-expressed with Pto/AvrPto in *NRC*-silenced (*NRC2a/b* and *NRC3*) *N. benthamiana*. Infiltrations without Pto/AvrPto or without NRC homologs were used as controls. HR index was established at 7 days post infiltration (dpi). Bars represent mean ± SE of 16 infiltrations from one biological repeat. Statistical differences among the samples were analysed by Student’s *t*-test (\**p*-value < 0.01). Experiments were performed at least three times with similar results. **(B)** Overexpression of Sl-NRC3 enhances Pto- and Cf-4- mediated cell death. Sl-NRC1, Sl-NRC2 and Sl-NRC3 were co-expressed with different immune receptors and AVR proteins, or autoactive immune receptor (Mi-1.2^T557S^), in leaves of *N. benthamiana*. Infiltrations without immune receptor were used as negative controls. HR index was established at 7 days post infiltration (dpi). Bars represent mean ± SE of 24 infiltrations from one biological repeat. Statistical differences among the samples were analysed by Student’s *t*-test (\**p*-value < 0.01). Experiments were performed at least three times with similar results.

